# ΔNP63 defines an exocrine-committed multipotent progenitor subset in the murine pancreas

**DOI:** 10.1101/2024.12.10.627728

**Authors:** Katarina Coolens, Melissa Van der Vliet, Jente Van Campenhout, Natalia del Pozo, Alejo Torres-Cano, Catharina Olson, Jianming Xu, Heiko Lickert, Meritxell Rovira, Isabelle Houbracken, Jonathan Baldan, Francisco X. Real, Francesca M. Spagnoli, Ilse Rooman

## Abstract

Cellular plasticity underpins heterogeneity in embryogenic progenitor cells and cancer cells. The transcription factor *deltaNp63* (*ΔNp63*) has been implicated in regulating cellular plasticity in several epithelial tissues. Despite a recently established role in steering plasticity of pancreatic cancer, ΔNp63 remains unstudied in pancreatic development.

Using murine single-cell sequencing data and RNA and protein *in situ* stainings, we assessed the spatio-temporal expression of *Trp63* and ΔNP63 in the embryonic pancreas. ΔNP63 demonstrates a transient and spatially restricted expression in the multipotent pancreatic progenitor (MPP) compartment delineating pro-exocrine progenitor cells. Lineage tracing of TP63^+^ cells marks a subset of MPPs and descendant exocrine acinar and centro-acinar/terminal duct cells. Lack of ΔNP63 in knock-out mice leads to hypotrophic exocrine acini with reduced levels of differentiation markers.

In summary, ΔNp63 confers heterogeneity within the MPP compartment, supporting exocrine cell development. These new insights in developmental plasticity have potential implications for pancreatic regeneration and cancer.

## Introduction

Cellular plasticity refers to the ability of cells to alter their fates or adopt entirely new identities^1^. Hence, such plasticity can contribute to heterogeneity of cell populations. During embryogenesis, multipotent progenitor cells possess inherent plasticity and recent single-cell studies have unveiled heterogeneity among embryonic progenitor cells in several tissues^2,3^. As development advances and cells undergo specialization, their plasticity progressively diminishes^4^. It remains important to decipher the regulators of such lineage restriction. Moreover, certain cells retain a limited degree of plasticity into adulthood, which underpins their capacity to contribute to tissue repair and regeneration^5^ and reinstitution of cellular plasticity has recently been recognized as a hallmark of cancer^6^. A more profound understanding of embryonic progenitor cell plasticity and how it is progressively restricted may therefore yield essential insights into the mechanisms that regulate regeneration and cancer.

The transcription factor deltaNp63 (*ΔNp63*), an isoform of the *Trp63* gene and a member of the *Trp53* family of transcription factors^7,8^, has been described as a key regulator of cellular plasticity in pancreatic ductal adenocarcinoma (PDAC) where it drives the most aggressive subtype^9^. Furthermore, we identified rare ΔNP63^+^ cells in the adult human pancreatic ductal system with higher occurrence during chronic pancreatitis, a condition where the tissue may regenerate but also can predispose to PDAC^10^. ΔNP63 was originally identified as a marker for multipotent organ-specific adult stem cells in various tissues^11–14^ and ΔNP63^+^ cells have been reported in developing lung and airways, mammary gland, skin and others^12,15–21^. To date, expression of ΔNP63 in the embryonic pancreas has not been studied.

Pancreatic embryonic development starts with the specification of multipotent pancreatic progenitors (MPP) in the dorsal and ventral anlagen of the posterior foregut endoderm at around embryonic day 9-9.5 (E9-9.5)^22^. Specified pancreatic progenitors, marked by PDX1, SOX9, HNF1B and PTF1A, expand the pancreatic anlagen that will form bud-like structures and subsequently, due to gut rotation, the two pancreatic buds will fuse together^23,24^. At E12-E12.5 the pancreatic epithelium further expands and undergoes branching morphogenesis, which starts with the formation of primary branches and concurrent regionalization into tip and trunk domains^25^. Progenitor cells in the tip domain express the marker CPA1 and remain multipotent^26^, yielding acinar and trunk cells, until E13.5-E14.5, becoming then restricted to the acinar cell lineage^26,27^. Recently, heterogeneity has been underscored within the MPP compartment, highlighting the existence of different progenitor populations^28,29^. Trunk progenitor cells are bipotent giving rise to both ductal and endocrine lineages^30^. The adult pancreas eventually consists of endocrine islets scattered throughout the exocrine acinar tissue which produces digestive enzymes (amylase, elastase, etc.) drained into the exocrine pancreatic ductal system, with centroacinar-terminal duct (CA-TD) cells at the interface between acinar cells and the ductal epithelium in the pancreas^31^.

We investigated the role of ΔNP63 during embryonic pancreas development by utilizing murine single-cell sequencing data, ΔNP63-lineage tracing mice, and a ΔNP63 knockout mouse model. Our findings reveal a transient expression and a critical role of ΔNP63 in the MPP compartment. Our study sheds light on the heterogeneity of the MPPs and the regulation of cell lineage commitment during pancreas organogenesis while also expanding the currently very limited knowledge on ΔNP63 in the pancreas.

## Results

### ΔNP63 displays a dynamic spatio-temporal expression pattern in the embryonic pancreatic exocrine tissue

To investigate the expression of *Trp63* during pancreatic development, we first re-analysed published sc-RNASeq (single cell RNA sequencing) datasets from murine pancreas from E10.5 (GSE144103), E12.5, E14.5 and E17.5 (GSE101099) (Figure 1a-f and Supplementary Fig. 1). We found a dynamic spatio-temporal expression pattern of *Trp63*, being enriched in the pancreas at E10.5 (Embryonic day 10.5) (Figure 1a,b), and then from E12.5 on becoming restricted to the epithelial tip (E-Tip) compartment and its descendants (Figure 1c-f). Specifically, we performed subclustering of all epithelial cells (denoted as "E") based on E-cadherin expression and annotated the clusters using specific markers. At E12.5, markers included *Cpa1*, *Ptf1a*, and *Bhlha15* for the E-Tip cluster; *Spp1* for the E-Trunk cluster; *Neurog3* and *Gcg* for the Endocrine cluster; and *Top2a*/*Mki67* for proliferating cells (Supplementary Fig. 1a-h). At E14.5 and E17.5, additional markers were used, including *Amy1* for the acinar cluster, *Try10* for maturation, and *Spp1* for the ductal cluster (Supplementary Fig. 1i-v). Notably, Trp63 was expressed in both proliferating cells (*Top2a/Mki67*-expressing cells) at E12.5 (2.72%) and E14.5 (2.30%), as well as in non-proliferating tip/acinar cell compartments (1.60% at E12.5 and 1.64% at E14.5) (Figure 1c-f). By E17.5, *Trp63* expression became restricted to very few acinar cells (0.15% of acinar cells, Supplementary Fig. 1w,x), which is in line with the low expression retained in adult murine pancreas^10^ (GSE109774^32^; Supplementary Fig. 2a-h). Additionally, *Trp63* was rarely detected in the trunk and ductal compartments at E12.5 (0.09%) and E14.5 (0.18%). Since the proportion of *Trp63*-expressing cells was the highest at E12.5, we performed a differential gene expression analysis at this stage between *Trp63*^+^ tip cells (red) and *Trp63*^-^ tip cells (blue) to generate the expression profile of *Trp63*-expressing cells (Supplementary Fig. 2i,j). This highlighted 42 significantly upregulated genes and no significantly downregulated genes (Adjusted p-value ≤ 0.05; Willcoxon Rank Sum Test) (Supplementary Table 1). Upregulated genes included acinar enzymes and acinar specific transcription factors, such as *Cpa2*, *Cel*, *Ctrb1*, *Pnliprp1*, *Cela1* and *Xbp1* (Figure 1g), and were identified as “protein digestion and absorption”, “pancreas secretion” and “fat digestion and absorption” by gene ontology analysis (GO terms) (Figure 1h).

**Figure 1:**
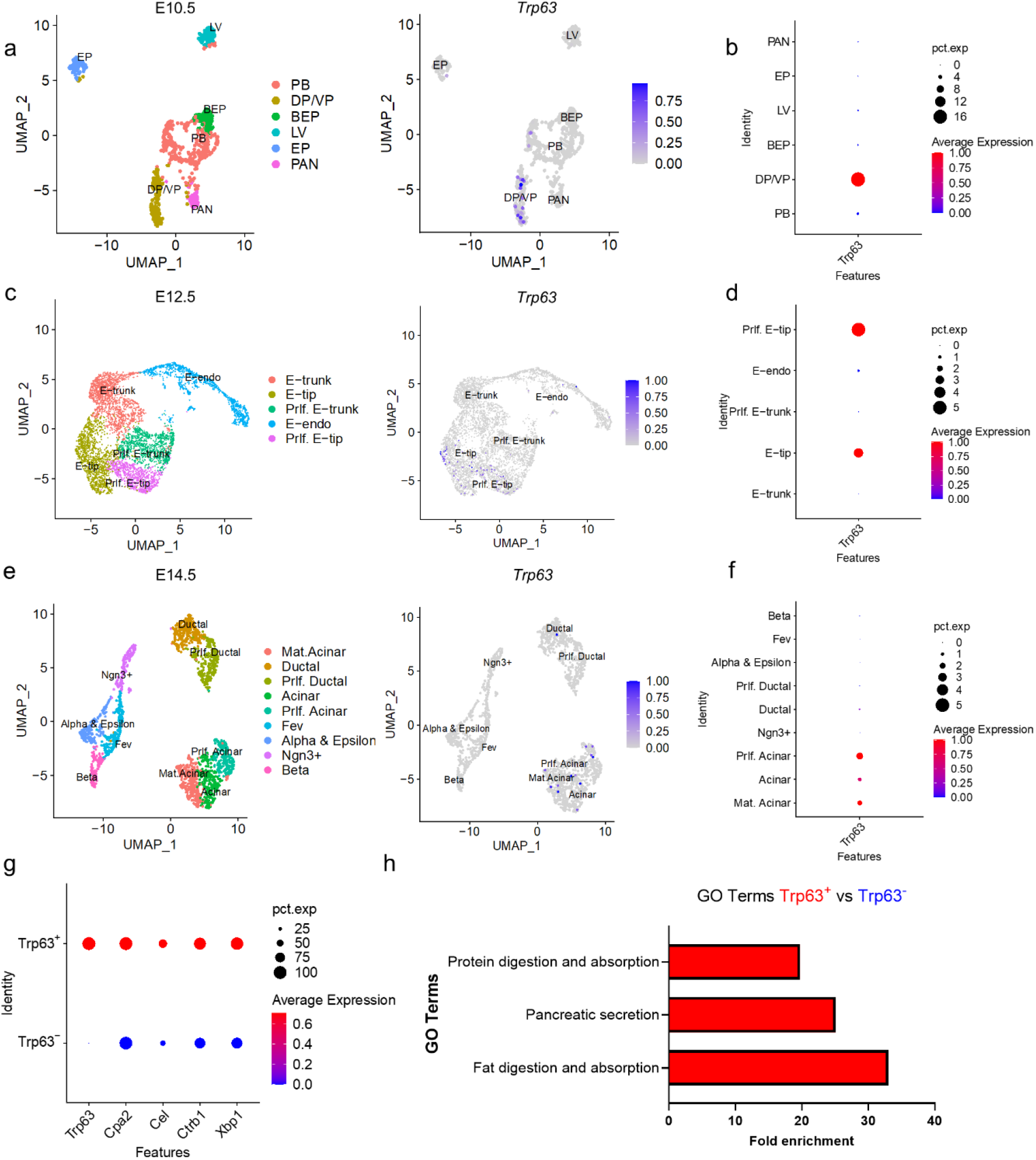
*Trp63* expression during embryonic development of the pancreas. **a-f** Re-analysis of single-cell transcriptomics data sets (GSE144103 and GSE101099) from murine embryonic pancreata at indicated timepoints. UMAP cell-cluster representation (left panels) and expression of *Trp63* as featureplots (middle panels) and as dotplots (right panels) at **a,b** E10.5 (*n*=3 mice), **c,d** E12.5 (*n*=2 mice) and **e,f** E14.5 (*n*=2 mice). **g** Dotplot of differential gene expression analysis between *Trp63-*positive (*Trp63*^+^) and *Trp63-*negative (*Trp63*^-^) tip cells showing *Trp63*, and the pancreatic digestive enzymes *Cpa2*, *Cel*, *Ctrb1*, *Cela1* and *Pnliprp1*. Adjusted p-value ≤ 0.05; Willcoxon Rank Sum Test. **h** Fold Enrichment in *Trp63-*positive (*Trp63*^+^) and *Trp63-*negative (*Trp63*^-^) cells for GO terms capturing the upregulated genes resulting from differentially expressed gene analysis based on panel g, showing enrichment in “Protein digestion and absorption”, “Pancreas secretion” and “Fat digestion and absorption”. UMAP = Uniform Manifold Approximation and Projection; PB = Pancreato-Biliary; DP/VP = Dorsal Pancreatic/Ventral Pancreatic bud; BEP = Bipotent Endoderm Progenitor; LV =Liver; PAN = PANcreatic; E = Epithelial. pct.exp = percentage expressed.

These observations prompted us to resolve stage-specific (E10.5, E12.5, E14.5 and E17.5) protein expression with specific focus on the ΔNP63 isoform. We employed an anti-TP63 antibody recognizing both ΔNP63 and TAP63 isoforms (Supplementary Fig. 3a). In addition, we used two distinct ΔNP63-specific antibodies (Figure 2), one of which we previously validated by Western blotting^10^ and that also stained the developing skin tissue (Supplementary Fig. 4). We confirmed our observations using a *ΔNp63*-specific BaseScope probe (Supplementary Fig. 3b).

**Figure 2:**
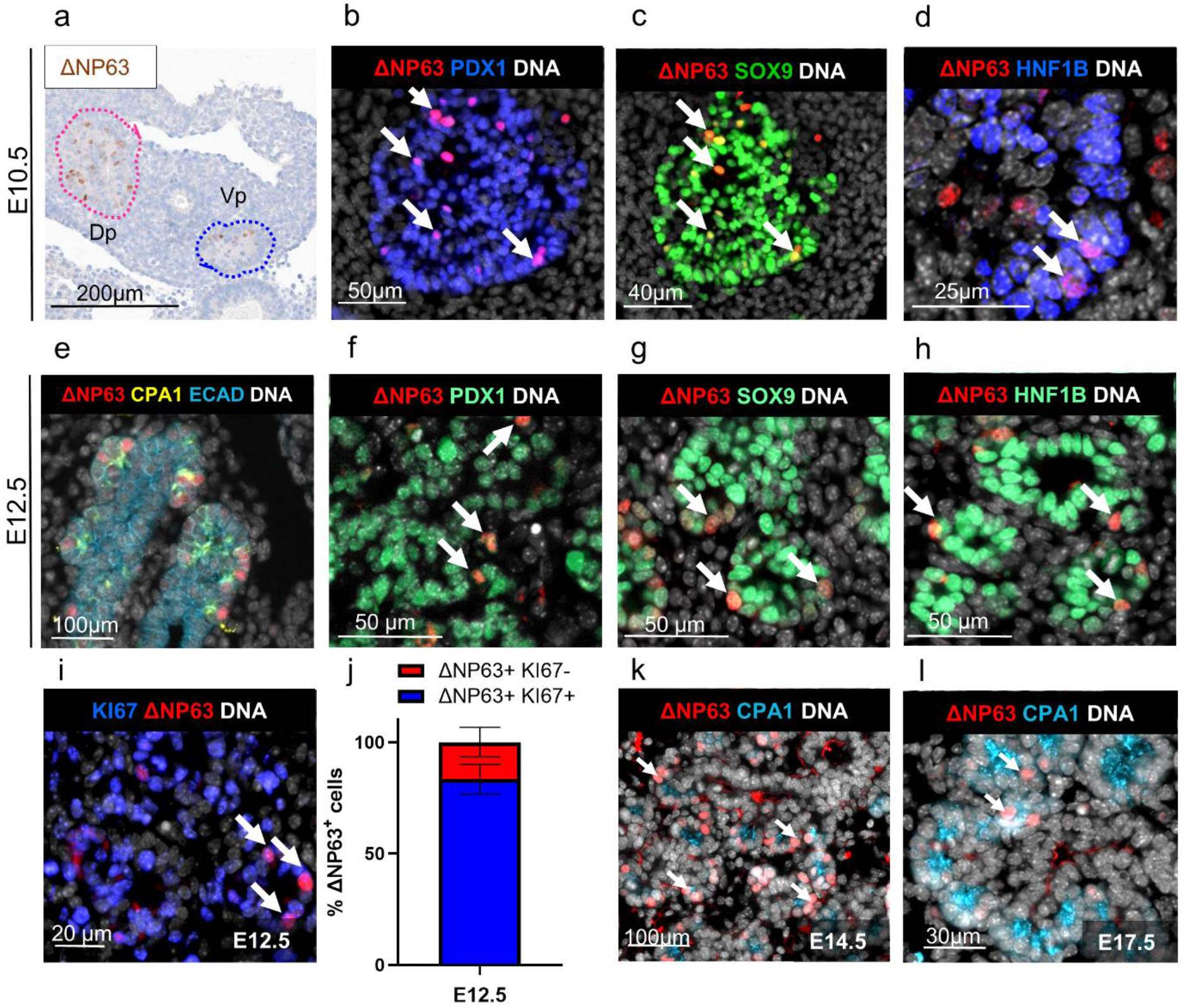
ΔNP63 protein expression in the developing murine pancreas. **a** Immunohistochemical staining of ΔNP63 in E10.5 dorsal (Dp) and ventral pancreatic (Vp) bud. **b** Immunofluorescence staining for ΔNP63 (red), DNA (white) and PDX1 (blue) at E10.5. **c** Immunofluorescence staining for ΔNP63 (red), DNA (white) and SOX9 (blue) at E10.5 **d** Immunofluorescence staining for ΔNP63 (red), DNA (white) and HNF1B (blue) at E10.5. **e** Immunofluorescence staining for ΔNP63 (red), DNA (white), CPA1 (yellow) and ECAD (blue) at E12.5. Immunofluorescence staining for ΔNP63 (red), DNA (white), **f** PDX1 (green), **g** SOX9 (green) or **h** HNF1B (green) at E12.5. **i** Immunofluorescence staining for ΔNP63 (red), DNA (white) and KI67 (blue) at E12.5. **j** Percentage of ΔNP63^+^ cells positive for KI67 (red bar) and negative for KI67 (blue bar) at E12.5; results are represented as mean ± SEM. Immunofluorescence staining for ΔNP63 (red), DNA (white) and CPA1 (blue) at **k** E14.5 and **l** E17.5.

At E10.5, we detected ΔNp63 in both pancreatic buds (Figure 2a, Supplementary Fig. 5a). The majority of ΔNP63^+^ cells (71.9 ± 4.8%) was located within an ECAD^high^ compartment and a smaller fraction (28.1 ± 4.8%) within an ECAD^low^ compartment that corresponds to the early pool of Glucagon^+^ endocrine progenitors (Supplementary Fig. 5b-e). In the ECAD^high^ compartment, ΔNp63^+^ cells were positive for typical markers of pancreatic MPPs (*i.e.,* PDX1, SOX9, HNF1B) (Figure 2b-d, Supplementary Fig. 5f). At E12.5, ΔNP63 protein was detected at the distal tips of the epithelial branches, marking 10.2 ± 0.3 % of CPA1^+^ cells (Figure 2e, Supplementary Fig. 5a, g), which is comparable to the 7% of all tip cells expressing *ΔNp63* in the sc-RNAseq (Figure 1d). Additionally, ΔNP63^+^ cells in the tip expressed MPP markers (SOX9, HNF1B and PDX1) (Figure 2f-h), the acinar transcription factor MIST1 (Supplementary Fig. 5h) and the majority (83.3 ± 3.8 %) of them were actively proliferating, as shown by the KI-67 staining (Figure 2i,j). ΔNP63 continued to mark a subset of the CPA1^+^, MIST1^+^ acinar progenitors until further reduction at E17.5 (Figure 2k,l and Supplementary Fig. 5i,j). Overall, our results revealed a consistent trend of ΔNP63 protein with that observed of *Trp63* in the single cell data (Figure 1g), showing the highest expression of ΔNP63 at embryonic day 12.5 (6.79 ± 0.35% of ECAD+ cells) followed by a progressive decrease (4.27 ± 0.29% at E14.5 and 0.26 ± 0.04% at E17.5) (Supplementary Fig. 5k).

In summary, ΔNP63 exhibits a transient expression pattern defining a subset of MPP, i.e. the pro-acinar cell lineage, with confinement to acinar cells at later stages.

### Descendants of TP63^+^ MPPs contribute to the exocrine compartment

To investigate the cell type specific contribution of the TP63/ΔNP63^+^ progenitors, we performed lineage analysis using *Trp63*-CreERT2; R26-mTmG mice^14^. The membrane localized tomato (mT) is constitutively expressed in all cells, after 4-hydroxytamoxifen (4-OHT) administration the expression of mT is replaced by membrane-localized GFP (mGFP) in *Trp63*-expressing cells and their descendants (Figure 3a). When 4-OHT was administered at E12.5 and mGFP^+^ cell distribution was assessed at E14.5 and E18.5 (Figure 3b), rare mGFP^+^ cells were detected at E14.5 (Figure 3c,d), alongside a non-induced control that confirmed the absence of leakiness of the reporter model (Supplementary Fig. 6a). mGFP^+^ cells exhibited markers of MPP’s, including CPA1 and PDX1 (Figure 3c, Supplementary Fig. 6b), as well as of acinar fate (MIST1) (Figure 3d, Supplementary Fig. 6c). At E18.5, mGFP^+^ *Trp63*-descendant cells (Figure 3e) belonged to the mature acinar cell lineage, as evidenced by co-staining with CPA1 and Amylase (AMY) (Figure 3e area 1, 2 and 3, Supplementary Fig. 7a-c). Interestingly, mGFP^+^ TP63 descendants were also observed in the center of the acinus, in the position of centroacinar cells (CA), and in KRT7^+^ terminal ductal cells (TD)^33^ (Figure 3 area 4 and 5, Supplementary Fig. 7d,e). Of all mGFP+ cells, 96.4 ± 1.6% could be detected in the acinar cells and 3.6 ± 1.6% in the CA-TD compartment (Supplementary Fig. 7f).

**Figure 3:**
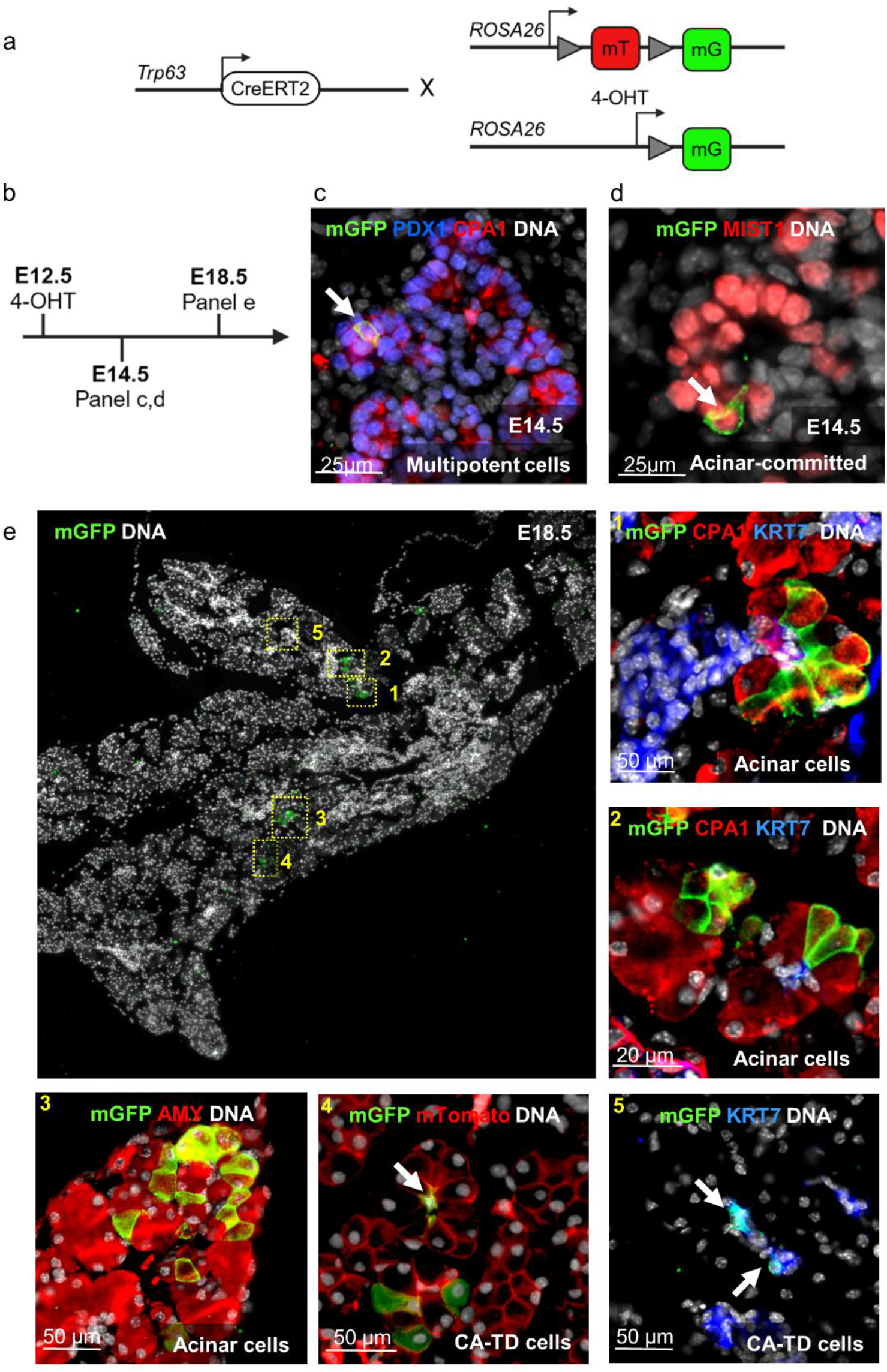
*Trp63* lineage tracing labels multipotent progenitors of the developing pancreas and later becomes lineage-restricted to (centro-)acinar cells. **a** Schematic representation of the lineage tracing mouse model using a two-color fluorescent Cre-reporter allele with tdTomato (mT) expressed prior to excision and enhanced green fluorescent protein (mG) after excision. 4-OHT = 4 hydroxytamoxifen. **b** Schematic representation of 4-OHT administration and time points of analysis. **c** Immunofluorescence staining for mGFP (green and indicated by white arrow), CPA1 (red), PDX1 (blue) and DNA (white) at E14.5. **d** Immunofluorescence staining for mGFP (green and indicated by white arrow), MIST1 (red) and DNA (white) at E14.5. **e** Overview of the analyzed tissue and numbered regions at higher magnification: **Region 1** Immunofluorescence for mGFP (green), CPA1 (red), KRT7 (blue) and DNA (white) at E18.5. **Region 2** Immunofluorescence for mGFP (green), CPA1 (red), KRT7 (blue) and DNA (white) at E18.5. **Region 3** Immunofluorescence for mGFP (green), AMY (red), and DNA (white) at E18.5. **Region 4** Immunofluorescence for mGFP (green), mTomato (red) and DNA (white) at E18.5. **Region 5** Immunofluorescence for mGFP (green), KRT7 (blue) and DNA (white) at E18.5. CA-TD = Centro-Acinar Terminal Ductal cells.

Our results demonstrated that TP63^+^ cells labelled at E12.5 give rise to acinar and CA-TD cells.

### Lack of ΔNP63 results in hypotrophic exocrine acini with impaired differentiation

Having established the dynamic, stage-dependent distribution of ΔNP63^+^ cells during pancreas development, we set out to determine whether *ΔNp63* regulated the organ development using a *ΔNp63^GFP^*^/+^ knock-in mouse strain, in which the *ΔNp63* coding sequence is replaced by *GFP*^34^. *ΔNp63^GFP/GFP^* homozygous embryos were stained for ΔNP63 and GFP (Figure 4a,b). 3D rendering showed a microscopic difference in some pancreatic lobes in knock-out (KO) mice at E14.5 with less dense tissue (red boxes in Figure 4c). Quantitative analysis of pancreatic weight and length at E16.5 underscored volume reduction of the pancreas (Figure 4d-f). Additionally, the affected lobes were also identifiable at the histological level showing more inter-acinar space and a disorganized pattern(Figure 4g). To understand what preceded this phenotypic difference, bulk RNAseq analysis of KO versus wild type pancreata at E12.5 was done. It highlighted 4372 differentially expressed genes (Adjusted p-value ≤ 0.05, Wald Test with Benjamini-Hochberg correction; Supplementary Table 2). Of these, 2299 genes were downregulated and another 2073 were upregulated. In addition to *Trp63*, the downregulated genes included the pro-acinar transcription factors *Mecom*^35^(Fold Change (FC) = -2.12) and *Gata4*^36^ (FC = -1.59), and early expressed acinar digestive enzymes such as *Ctrb1*^37^ (FC = -7.1) and *Cela1*^38^ (FC = -1.59) (Figure 5a). Consistently, gene set enrichment analysis (GSEA) showed that gene sets significantly downregulated in ΔNP63 KO mice belonged to the ‘Exocrine Pancreas development’ GO term (Figure 5b). Upregulated genes featured pro-endocrine genes, including *Fev* (FC = 1.64), *Pdx1* (FC = 1.97) and *Chga* (FC = 2.15) (Figure 5c) and were enriched in GO term “Insulin secretion” (Figure 5d) next to *Pdx1*, *Sox9* (FC = 0.91) and *Hnf1b* (FC = 1.2), markers of MPPs at that stage of development (Figure 5c). In line with the transcriptome analysis, KO pancreata showed fewer CPA1^+^ tip cells (Figure 5e). Later, at E16.5, the acini were immature with diminished levels of Amylase and E-cadherin^39^ (Figure 5f and Supplementary Fig. 8a), as well as reduced expression of other acinar enzymes and transcription factors (Figure 5g). Additionally, we measured acinus diameter, knowing that immature acinar cells are typically distinguished by their smaller volume^40^. Indeed, acinar cells in the KO mice demonstrated a significant reduction in cell diameter (Figure 5h). KO pancreata exhibited no significant differences in cell proliferation compared to WT mice, including at earlier developmental stages (Figure 5i and Supplementary Fig. 8b-d). Additionally, only a minimal number of apoptotic cells were observed in the pancreatic epithelium, with no discernible differences between WT and KO samples (Supplementary Fig. 8b, lower panel). At this timepoint, endocrine and ductal markers were not affected (Supplementary Fig. 8e), indicating no differences in lineage commitment but in the differentiation state of acinar cells that remains more immature.

**Figure 4:**
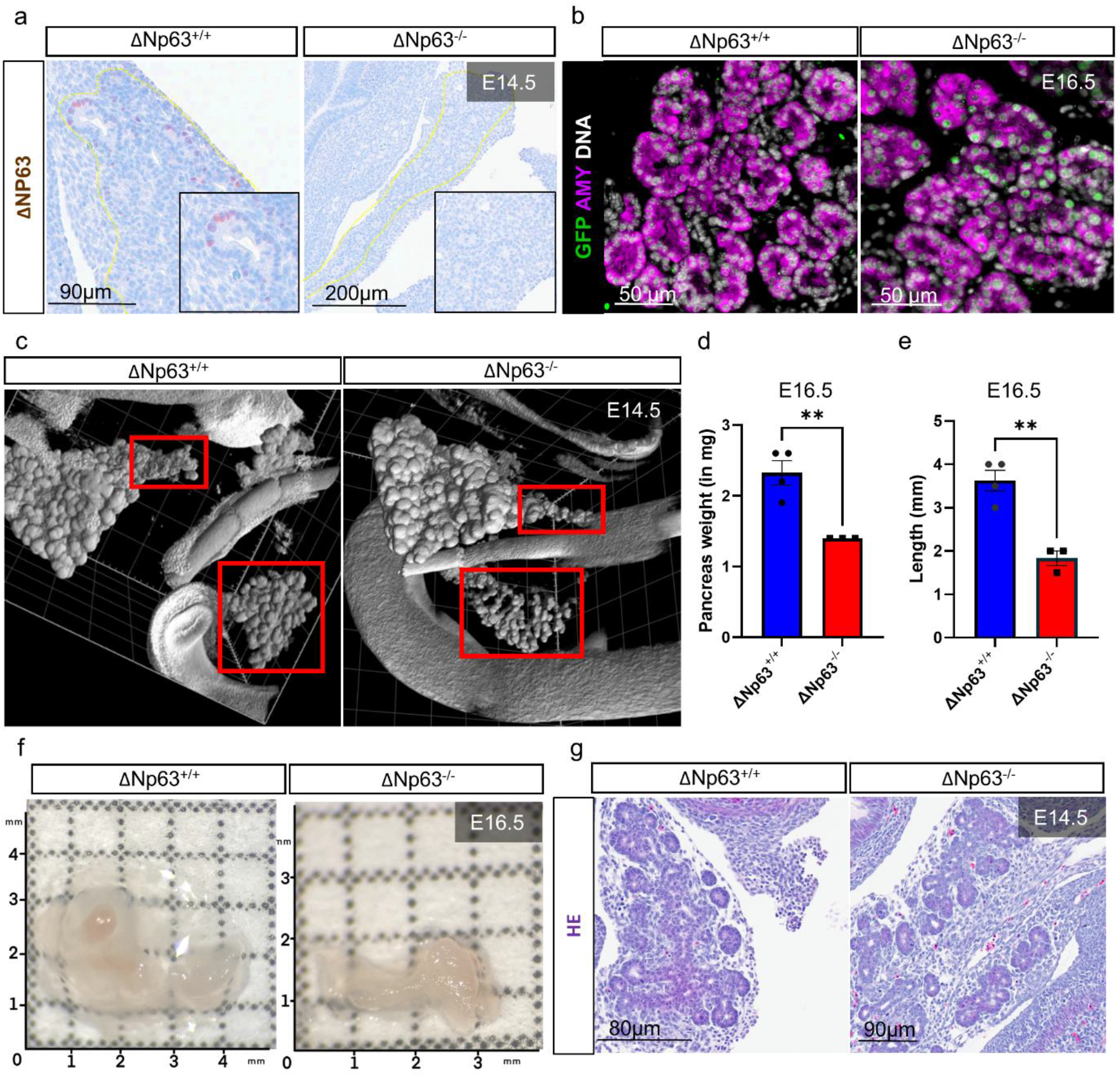
Pancreatic phenotype in ΔNP63 knockout mice. **a** Immunohistochemical staining of ΔNP63 in pancreas at E14.5 in ΔNP63 wildtype (WT, ΔNp63^+/+^) and knockout (KO, ΔNp63^-/-^) mice. **b** Immunofluorescence for GFP (green), AMY (purple) and DNA (white) in ΔNP63 wildtype (WT, ΔNp63^+/+^) and knockout (KO, ΔNp63^-/-^) mice at E16.5. **c** 3D rendering of the gastrointestinal tract of ΔNp63 wildtype (WT, ΔNp63^+/+^) and knockout (KO, ΔNp63^-/-^) mice at E14.5. Red boxes highlight the most affected regions of the pancreas, being the caudal part of the splenic lobe and the ventral lobe. **d** Quantification of weight (mg) in ΔNP63 wildtype (WT, ΔNp63^+/+^) and knockout (KO, ΔNp63^-/-^) mice at E16.5; **p ≤ 0.01; Unpaired two-tailed t-test was performed; *n*=3 mice were analyzed. **e** Quantification of length (mm) in ΔNP63 wildtype (WT, ΔNp63^+/+^) and knockout (KO, ΔNp63^-/-^) mice at E16.5; **p ≤ 0.01; Unpaired two-tailed t-test was performed; *n*=3 mice were analyzed. **f** Example of extracted embryonic pancreata of wildtype (WT, ΔNp63^+/+^) and knockout mice (KO, ΔNp63^-/-^) at E16.5. **g** Haematoxylin-eosin staining of the pancreata of wildtype (WT, ΔNp63^+/+^) and knockout (KO, ΔNp63^-/-^) mice at E14.5.

**Figure 5:**
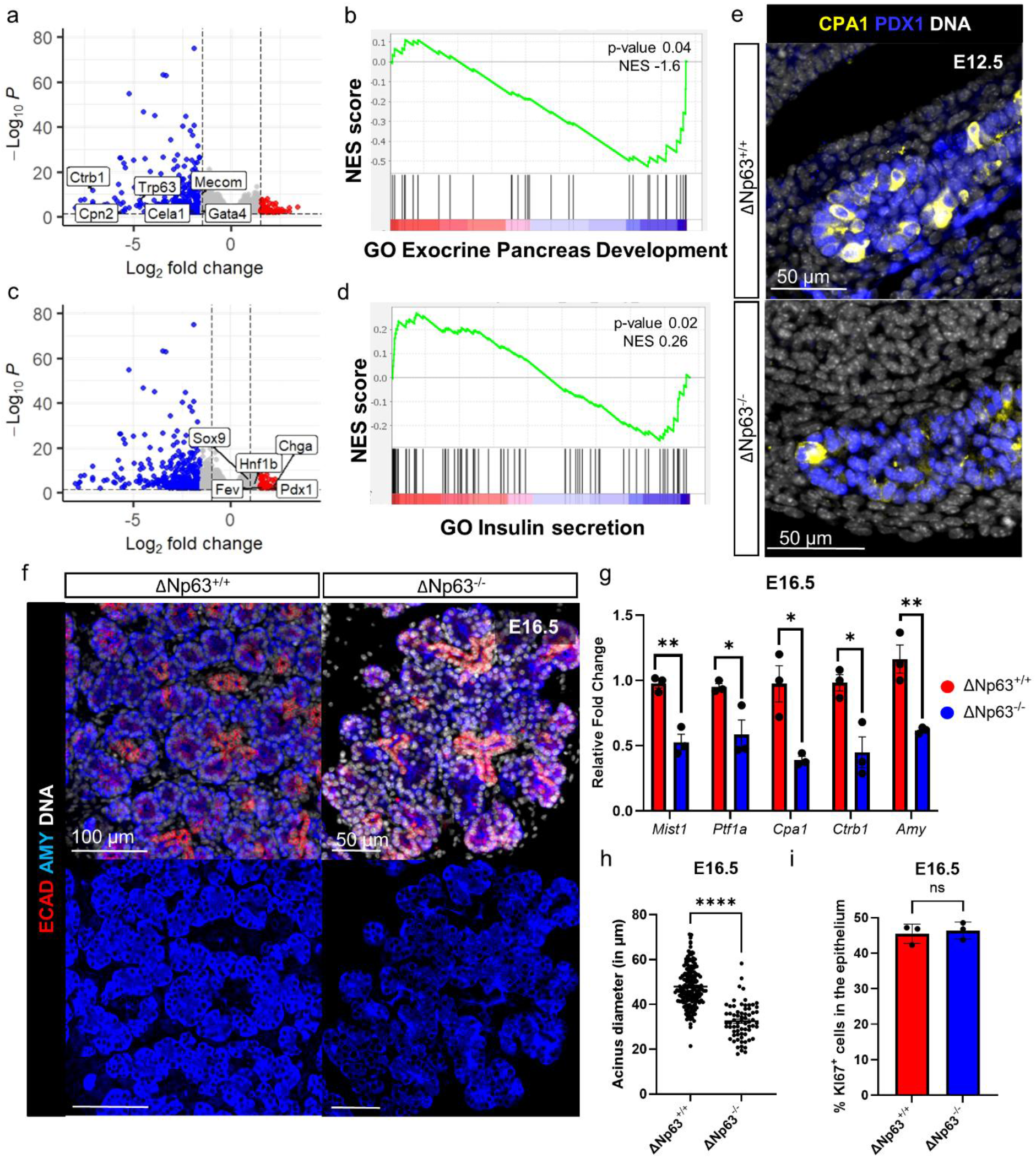
Affection of the acinar compartment in ΔNP63 knockout mice. **a** Volcano plot highlighting the downregulated genes of interest from the differentially gene expression analysis between ΔNP63 knockout (KO, ΔNp63^-/-^) and wildtype (WT, ΔNp63^+/+^) mice at E12.5; Adjusted p-value ≤ 0.05, Wald Test with Benjamini-Hochberg correction was performed. **b** Gene Set enrichment Analysis (GSEA) of the downregulated genes from panel a. **c** Volcano plot highlighting upregulated genes of interest from the differentially gene expression analysis between ΔNP63 knockout (KO, ΔNp63^-/-^) and wildtype (WT, ΔNp63^+/+^) mice at E12.5; Adjusted p-value ≤ 0.05, Wald Test with Benjamini-Hochberg correction was performed. **d** GSEA analysis of the upregulated genes from panel c. **e** Immunofluorescence for CPA1 (yellow), PDX1 (blue) and DNA (white) in ΔNP63 wildtype (WT, ΔNp63^+/+^) and knockout (KO, ΔNp63^-/-^) mice at E12.5. **f** Immunofluorescence for ECAD (red), AMY (blue) and DNA (white) in ΔNP63 wildtype (WT, ΔNp63^+/+^) and knockout (KO, ΔNp63^-/-^) mice at E16.5. **g** Quantitative RT-PCR of exocrine-specific genes measure in ΔNP63 wildtype (WT, ΔNp63^+/+^) and knockout (KO, ΔNp63^-/-^) mice at E16.5 ( *p≤0.05 and **p≤0.01); Unpaired two-tailed t-test was performed; *n*=3 mice were analyzed. **h** Quantification of the diameter (μm) of the acinus in ΔNP63 wildtype (WT, ΔNp63^+/+^) and knockout (KO, ΔNp63^-/-^) mice at E16.5; ****p ≤ 0.0001; TWO-way anova for repeated measures was performed. **i** Quantification of the percentage of KI67^+^ cells in the epithelium (based on ECAD expression) in ΔNp63 wildtype (WT, ΔNp63^+/+^) and knockout (KO, ΔNp63^-/-^) mice at E16.5; *ns* = non-significant; Unpaired two-tailed t-test was performed; *n*=3 mice were analyzed.

In summary, lack of ΔNP63 in the developing pancreas results in hypotropic exocrine acini with reduced differentiation markers.

## Discussion

Biological principles often reproduce in multiple tissues. Hence, regulation of cellular plasticity as it occurs during embryonic development, tumor initiation and cancer progression in epithelia of lung, mammary gland, skin and other organs, may also apply to the pancreas. Following several studies on ΔNP63’s role in steering the cell fate in pancreatic cancer^9,41^, we reported the existence of rare ΔNP63^+^ cells in the adult human exocrine pancreas^10^. The transcription factor ΔNp63 has been extensively documented in aforementioned epithelial tissues where it is implicated in developmental processes and cancer ^12,16,18,20,42,43^. Here, we detected ΔNP63 as soon as the budding of the pancreatic epithelium is started, at around embryonic day E10.5, and its expression is extinguished by E17.5 when the pancreas already contains all mature endocrine and exocrine cell types. In early developing pancreas, ΔNP63 was confined to the developing tip cells (Figure 1 and 2). The pancreatic tip cell domain, marked by CPA1^26^, has long thought to be a very homogeneous domain, containing only cells that are equally multipotent. Our extraction of a *Trp63* expression profile out of the single cell sequencing showed that *Trp63^+^* tip cells are already more acinar committed at E12.5. For this we found support in the RNAseq data of Yu *et al*. where a significant downregulation of *Trp63* can be noted in the trunk cell compartment compared to the acinar compartment^23^. This is also in line with a clonal tracing analysis suggesting that acinar-primed progenitors might already be present in this multipotent cell population^27^. Our *in vivo* lineage tracing assays established the presence of TP63^+^ cells among both multipotent and acinar-committed cells, and their contribute to the establishment of mature (centro) acinar cells. Our results thus hint at the presence of (centro) acinar-committed tip cells, underscoring heterogeneity within the MPPs. Importantly, we also provide for the first time a marker specific of these acinar-committed tip cells amongst the multipotent ones.

The labelling efficiency in our study is low, which may be a limitation to reliably identify all descendant cells. Yet, it is consistent with other reports. For instance, CPA1 tracing, a more abundantly expressed marker compared to ΔNP63, showed a maximum tracing efficiency of 20%^26^. Similarly, TP63 tracing in the lung, using Trp63-CreERT mice, reported labelling efficiencies ranging from 6.7% to 35.7%, depending on the region^18^. Since the single cell data and the *in situ* stainings indicated that ΔNP63^+^ cells in the embryonic pancreas are less prevalent compared to skin for example, and that ΔNP63 is expressed at lower level, a lower tracing efficiency was expected. Despite labelling only few cells, these experiments showed contribution of TP63^+^ cells to acinar cells and also to CA-TD, a cell type that is enriched for pancreatic progenitor/stem cell markers (PTF1A, SOX9, SCA-1, SDF-1, C-MET, and Nestin)^31,44^ but that has been less studied than the other pancreatic cell types. Besides an origin within the tip cell domain^31^, the developmental origin of CA cells or critical transcription factors herein have not been established^45,46^. Here, we suggest that Trp63^+^ cell descendants comprise CA-TD cells, warranting further investigation. Without a more efficient reporter, we cannot draw firm conclusions on the endocrine cell compartment. Furthermore, up until now we don’t have any further explanation to how our findings in the embryonic pancreas relate to the ΔNP63^+^ cells we found in the adult human pancreas, where they reside within the ductal epithelium^10^.

At E10.5, we observed co-expression of ΔNp63 with glucagon, a finding consistent with Yu et al.’s data, and identifying the first wave of alpha cells^23,47,48^. In our tracing study we did not observe any contribution to the mature endocrine compartment. This could be attributed to the early alpha cells dying ^49,50^ or it could be that early alpha cells contribute to mature ones within the second wave of NGN3 at E14.5^51^, all hypotheses that are still under debate. As our tracing was performed from E12.5 onwards -after the first peak of NGN3-, this might explain why we do not observe any contribution of TP63 to this compartment. Earlier attempts starting from E9.5, E10.5, or E11.5 were unsuccessful, likely due to low ΔNp63 expression levels. Increasing the 5-OHT dose increased the risk of fetal loss.

While postnatal stages could not be analyzed because of the embryonically lethal phenotype^34^, we unveiled a role of ΔNP63 in acinar cell differentiation as the KO pancreata at E16.5 showed a decrease in acinar cell specific gene expression and more immature acini. At E12.5 we already noted an increase of MPP markers *Sox9*, *Hnf1b* and *Pdx1* potentially indicating that exhaustion of CPA1^+^ MPP cells triggered replenishment of early MPP’s to further support differentiation and development of the pancreas.

Our 3D renderings revealed that the most pronounced phenotype occurred in the ventral and caudal part of the splenic lobe of the pancreas, despite *Trp63* expression being present in both the ventral and dorsal pancreatic buds. This observation may suggest the existence of compensatory mechanisms within the dorsal bud or the existence of other progenitor cell populations. A potential (partly compensatory) involvement of the *TAp63* isoform of *Trp63* can be evaluated *in situ*, but, at least at E12.5 the overall *Trp63* expression is reduced. Hence, we are currently unable to provide an explanation for the lobe-specific phenotype.

Our findings indicated that ΔNp63 functions to segregate the MPP population and directs these cells toward acinar and CA-TD cell fates, thereby playing a critical role in restraining cellular plasticity during embryonic development. With acinar cell plasticity playing a role in the initiation of PDAC^52,53^, it should be investigated if ΔNp63 could be at play. However, thus far, we haven’t been able to identify ΔNp63 in adult mouse models^10^, limiting our ability to study this. On the other hand, new studies on ΔNP63’s upstream and down-stream regulators^54–58^, potential interaction partners^59,60^ as well as modulating agents^61,62^ could be explored in steering pancreatic cell fate decisions, including those relevant for restraining exocrine cell development and potentially promoting endocrine cell (re)generation.

In conclusion, this first report of ΔNp63 in the embryonic pancreas opens new avenues for research on cellular plasticity in pancreatic disease and regeneration.

## Material and methods

### Mouse strains and experiments

All mice experiments were approved by the Ethical Committee for Animal Experimentation at Vrije Universiteit Brussel (#22-637). All animal procedures performed at CNIO were approved by local and regional ethics committees [(Institutional Animal Care and Use Committee and Ethics Committee for Research and Animal Welfare, Instituto de Salud Carlos III) (CBA 09_2015_v2) and Comunidad Autónoma de Madrid (ES280790000186)] and performed according to the European Union guidelines. Noon of the day the vaginal plug was detected, was considered as embryonic day E0.5. The wild-type CD-1 females, used for characterization of the cells, were purchased from Janvier. For lineage tracing experiments, Trp63-CreERT2 mice^14^ were crossed with Rosa26-flox-mTRed-Stop-flox-mGFP mice (Jackson Laboratory #007576) at the Epithelial Carcinogenesis Group, Spanish National Cancer Research Centre. A dose of 5mg 4-hydroxytamoxifen (4-OHT, Sigma, Saint Louis, USA, H6278) was administrated at embryonic time point E12.5 and pregnant females were sacrificed at E14.5 and E18.5. 4-OHT was dissolved in sunflower seed oil (Sigma, C8267) and administered via intraperitoneal injection. ΔNp63^GFP^ KO mice were a kind gift of Prof. Romano (Buffalo University, department Oral Biology). KO mice were genotyped as previously described^34^.

### Single-cell RNA sequencing re-analysis

Single-cell expression patterns of *Trp63* were obtained from the publicly available datasets GSE144103 (E10.5)^28^ and GSE101099^63^, which contains murine embryonic pancreas samples at developmental stages E12.5, E14.5, and E17.5. GSE144103 was processed as previously described^28^. From GSE101099 the following samples were analyzed: GSM3140915 and GSM2699156 for E12.5, GSM2699154_E14_B1 and GSM2699155_E14_B2 for E14.5, and GSM3140917_E17_1_v2 and GSM3140918_E17_2_v2 for E17.5. Data preprocessing was performed using RStudio (version 4.4.0) with the "Seurat" package (version 5.1.0)^64^. To reduce noise, genes expressed in fewer than three cells and cells expressing fewer than 200 genes were excluded from all datasets. Additional quality control steps were taken to filter out low-quality cells, such as empty droplets. Specifically, cells expressing fewer than 500 genes were excluded from samples GSM2699156, GSM2699155_E14_B2, and GSM3140918_E17_2_v2, while cells expressing fewer than 600 genes were removed from samples GSM2699154_E14_B1 and GSM3140917_E17_1_v2. A more stringent threshold of 1300 genes was applied for GSM3140915. After filtering, samples from each developmental stage were merged, and the combined datasets were log-normalized. The top 2000 highly variable genes were identified for each timepoint using the variance-stabilizing transformation (VST) method. Standard Seurat workflows were subsequently applied to the datasets. Single-cell neighborhood graphs were constructed using principal component analysis (PCA), with the first 20, 19, and 21 principal components selected for E12.5, E14.5, and E17.5, respectively. Clustering was performed using the Louvain algorithm with a resolution of 0.1 for each timepoint. To account for batch effects, Harmony integration was applied when necessary. Additionally, the impact of cell cycle effects and mitochondrial gene expression was assessed, and these factors were regressed out during the data scaling process if deemed necessary. Cell cluster identities were determined through the use of specific marker genes, allowing for the identification of distinct cell types in the dataset. No new codes were generated for this analysis.

### Immunostainings

Embryonic tissue samples from different embryonic days were fixed overnight at room temperature in 4% paraformaldehyde solution (VWR, Pennsylvania, USA), followed by dehydration and embedding in paraffin. 4µm FFPE tissue sections were cut. For IHC-DAB stainings slides were baked for 1 hour at 60°C, followed by deparaffinization and rehydration. Endogenous peroxidase activity was blocked by using 3% Hydrogen Peroxidase in methanol for 30 minutes. Antigen retrieval was performed using citrate buffer (Sigma) and protein block was done using casein block (Thermofisher Scientific, Massachusetts, USA), diluted 1:4 in PBS. The primary antibody was incubated overnight at 4°C in a humidity chamber. The next day, slides were washed and incubated with the biotinylated secondary antibody for 30 minutes at room temperature. Next, the slides were washed and incubated for 5 minutes at room temperature with the Streptavidin-Biotin-HRP complex (VECTASTAIN® Elite® ABC kit (Standard)). Slides were washed again and were incubated with DAB and counterstained with haematoxylin. The list of antibodies can be found in Supplementary Table 3. For cell death detection, the *In Situ* Cell Death Detection Kit (11684795910, Merck, Darmstadt, Germany) was used.

For immunofluorescence staining of ΔNp63 (P40) the Alexa Fluor 647 anti-mouse Tyramide Superboost kit (Thermofisher Scientific) was used according to manufacturer’s protocol. Slides were washed with PBS-T and incubated overnight with other primary antibodies after the completion of the Superboost protocol for multiplex immunofluorescence stainings. The next day, slides were washed again and mounted with ProLong Gold Antifade Mountant (ThermoFisher Scientific).

### RNA in situ hybridization (BaseScope)

BaseScope RNA in situ hybridization was done on FFPE tissue according to manufacturer’s protocol with the BaseScope RED v2 kit (ACD, Biotechne, #323600). Probes for ΔNp63 (ACD Bio, #7147)41, PPIB (positive control, #712351) and Dapb (negative control, #701021) were purchased from Advanced Cell Diagnostics (Newark, USA).

### 3D staining: FLIP-IT

The gastrointestinal of both WT and KO embryos were collected in chilled PBS and were afterwards cleared and stained as previously described^10^. Samples were incubated with Hoechst (1:1000, ThermoFisher Scientific, # H1399).

### Imaging

DAB-IHC slides were visualized and scanned using the Leica GT-450. Fluorescent multiplex stainings were visualized with the EVOS FL Auto Cell Imaging System and were scanned using the Zeiss Axioscan Z.1. Post-processing was done using Zeiss Zenn Lite blue3.6. 3D samples were visualized using Zeiss Lighsheet Z.1 (Zeiss, Oberkochen, Germany). Images were acquired using 20x objective, NA=1 with zoom 0.36 – 8bit.

### Quantitative Reverse Transcription Polymerase Chain Reaction (qRT-PCR)

Total RNA from embryonic pancreata was isolated using the miRNeasy Micro Kit (Qiagen, Hilde, Germany). RNA concentration was measured using the NanoDrop 2000 (Thermofisher). Total RNA was reversed transcribed into cDNA using the GoScript Reverse Transcription System (Invitrogen). qPCR was performed using FastSYBRGreen 5× MasterMix on a QuantStudio 6 (Invitrogen). Primers were obtained from IDT (Supplementary Table 4). Analysis was done by determination of the comparative threshold cycle. For normalization GAPDH and HPRT were used.

### Bulk RNA-sequencing

The bulk RNA samples were sequenced on a NovaSeq 6000 system, hosted by BRIGHTcore (Brussels, Belgium). After demultiplexing with bcl-convert (version 3.7.5), the read quality was assessed with fastqc (http://www.bioinformatics.babraham.ac.uk/projects/fastqc/). The raw reads were mapped against the mouse genome (version GRCm38-83) using STAR^65^. The mapped reads were then translated into a quantitative measure of gene expression with the open-source tool (HTSeq)^66^. Following quantification of expression levels, we looked at the differential expression between the knock-out and wild type samples using the R/Bioconductor DESeq2 package^67^.

### Data analysis

The HALO image analysis platform (version v4.0.5107.318) was utilized for all quantifications of 2D slides. To calculate the percentage of cells expressing a certain marker we used the multiplex IHC v3.4 quantification algorithm.

### Statistics

Experimental data were analyzed using GraphPad Prism10.0 and statistical significance was accepted at p ≤ 0.05. The results are shown as mean ± standard error of mean (SEM). The datasets were assessed for normality using the Shapiro-Wilk test. Variables following a normal distribution were analyzed using the unpaired, two-tailed Student’s t-test and datasets that did not follow a normal distribution were analyzed using the Willcoxon Rank Sum Test. On the bulk RNA sequencing analysis data we performed the Wald Test with Benjamini-Hochberg correction, built in into the DESeq2 package. The number of independent experiments (n) is indicated in the figure legends. All IHC/IF/ISH assays were done on at least 3 biological replicates, at least 3 sections were analyzed for each replicate.

## Acknowledgments

We would like to thank the Visual and Spatial Tissue Analysis (VSTA) core facility. Sequencing was performed at BRIGHTcore (www.brightcore.be). We thank Catharina Olson for the analysis of the bulk RNA sequencing. In addition, we thank M. A. Blasco, E. Lapi, M. Ramal and the Epithelial Carcinogenesis Group members for their valuable contributions.

## Funding

This work is supported by Fonds Wetenschappelijk Onderzoek (FWO, G0A7322N) and Wetenschappelijk Fonds Willy Gepts. KC is a recipient of a PhD Fellowship of the FWO (1157221N). IH was financially supported by the bequests of Ms. Esther Desmedt and Ms. Irma Noë. JB is supported by a Foundation Against Cancer Fundamental mandate (2023-038). MR is supported by Ministerio de Ciencia, Innovación y Universidades (MICIN) (PID2022-136973OB-I00). Work in the laboratory of F.X.R. is supported by grant PID2021-128125OB-I00 from Ministerio de Ciencia, Innovación y Universidades (Madrid, Spain). CNIO is supported by MCIU as a Centro de Excelencia Severo Ochoa (grant SEV-2015-0510).

## Author contributions

This study was conceptualized and designed by KC and IR. KC, JVC and NDP performed experiments. MVDV performed all single cell sequencing re-analysis. KC performed data collection and interpretation. CO performed the analysis of the bulk RNA-sequencing. XJ generated the Trp63-CreERT2 mice. ATC, HL, MR, IH, JB, FXR and FS provided intellectual input. KC and IR wrote the manuscript, and all authors edited the manuscript.

## Funding

This work is supported by Fonds Wetenschappelijk Onderzoek (FWO, G0A7322N) and Wetenschappelijk Fonds Willy Gepts. KC is a recipient of a PhD Fellowship of the FWO (1157221N) and of the Award Cancer Research - Oncology Center Vrije Universiteit Brussel, funded by the bequests of Ms. Esther Desmedt and Ms. Irma Noë. IH was financially supported by the bequests of Ms. Esther Desmedt and Ms. Irma Noë. JB is supported by a Foundation Against Cancer Fundamental mandate (2023-038). MR is supported by Ministerio de Ciencia, Innovación y Universidades (MICIN) (PID2022-136973OB-I00). Work in the laboratory of F.X.R. is supported by grant PID2021-128125OB-I00 from Ministerio de Ciencia, Innovación y Universidades (Madrid, Spain). CNIO is supported by MCIU as a Centro de Excelencia Severo Ochoa (grant SEV-2015-0510).

## Conflict of interests

The authors declare that they have no conflict of interest.

## Materials & correspondence

Correspondence to Ilse Rooman

## References

1. Shen, S. & Clairambault, J. Cell plasticity in cancer cell populations. F1000Research 9, (2020).

2. Cui, Z., Wei, H., Goding, C. & Cui, R. Stem cell heterogeneity, plasticity, and regulation. Life Sci. 334, 122240 (2023).

3. Bedzhov, I., Graham, S. J. L., Leung, C. Y. & Zernicka-Goetz, M. Developmental plasticity, cell fate specification and morphogenesis in the early mouse embryo. Philos. Trans. R. Soc. London. Ser. B, Biol. Sci. 369, (2014).

4. Jahagirdar, B. N. & Verfaillie, C. M. Multipotent adult progenitor cell and stem cell plasticity. Stem Cell Rev. 1, 53–59 (2005).

5. Poulsom, R., Alison, M. R., Forbes, S. J. & Wright, N. A. Adult stem cell plasticity. J. Pathol. 197, 441–456 (2002).

6. Yuan, S., Norgard, R. J. & Stanger, B. Z. Cellular Plasticity in Cancer. Cancer Discov. 9, 837–851 (2019).

7. Mangiulli, M. et al. Identification and functional characterization of two new transcriptional variants of the human p63 gene. Nucleic Acids Res. 37, 6092–6104 (2009).

8. Fisher, M. L., Balinth, S. & Mills, A. A. p63-related signaling at a glance. J. Cell Sci. 133, (2020).

9. Somerville, T. D. D. et al. TP63-Mediated Enhancer Reprogramming Drives the Squamous Subtype of Pancreatic Ductal Adenocarcinoma. Cell Rep. 25, 1741–1755.e7 (2018).

10. Martens, S. et al. Discovery and 3D imaging of a novel ΔNp63-expressing basal cell type in human pancreatic ducts with implications in disease. Gut gutjnl-2020-322874 (2021) doi:10.1136/gutjnl-2020-322874.

11. Truong, A. B., Kretz, M., Ridky, T. W., Kimmel, R. & Khavari, P. A. p63 regulates proliferation and differentiation of developmentally mature keratinocytes. Genes Dev. 20, 3185–3197 (2006).

12. Signoretti, S. et al. p63 is a prostate basal cell marker and is required for prostate development. Am. J. Pathol. 157, 1769–1775 (2000).

13. Rock, J. R. et al. Basal cells as stem cells of the mouse trachea and human airway epithelium. Proc. Natl. Acad. Sci. 106, 12771–12775 (2009).

14. Lee, D.-K., Liu, Y., Liao, L., Wang, F. & Xu, J. The prostate basal cell (BC) heterogeneity and the p63-positive BC differentiation spectrum in mice. Int. J. Biol. Sci. 10, 1007–1017 (2014).

15. Pignon, J.-C. et al. p63-expressing cells are the stem cells of developing prostate, bladder, and colorectal epithelia. Proc. Natl. Acad. Sci. U. S. A. 110, 8105–8110 (2013).

16. Chakrabarti, R. et al. ΔNp63 promotes stem cell activity in mammary gland development and basal-like breast cancer by enhancing Fzd7 expression and Wnt signalling. Nat. Cell Biol. 16, 1–13,1004-1015 (2014).

17. Blanpain, C. & Fuchs, E. P63: Revving up epithelial stem-cell potential. Nat. Cell Biol. 9, 731–733 (2007).

18. Yang, Y. et al. Spatial-Temporal Lineage Restrictions of Embryonic p63(+) Progenitors Establish Distinct Stem Cell Pools in Adult Airways. Dev. Cell 44, 752–761.e4 (2018).

19. Yang, A. et al. p63 is essential for regenerative proliferation in limb, craniofacial and epithelial development. Nature 398, 714–718 (1999).

20. Mills, A. A. et al. p63 is a p53 homologue required for limb and epidermal morphogenesis. Nature 398, 708–713 (1999).

21. Karni-Schmidt, O. et al. Distinct expression profiles of p63 variants during urothelial development and bladder cancer progression. Am. J. Pathol. 178, 1350–1360 (2011).

22. Villasenor, A., Chong, D. C., Henkemeyer, M. & Cleaver, O. Epithelial dynamics of pancreatic branching morphogenesis. Development 137, 4295–4305 (2010).

23. Yu, X.-X. et al. Defining multistep cell fate decision pathways during pancreatic development at single-cell resolution. EMBO J. 38, (2019).

24. Burlison, J. S., Long, Q., Fujitani, Y., Wright, C. V. E. & Magnuson, M. A. Pdx-1 and Ptf1a concurrently determine fate specification of pancreatic multipotent progenitor cells. Dev. Biol. 316, 74–86 (2008).

25. Pan, F. C. & Wright, C. Pancreas organogenesis: from bud to plexus to gland. Dev. Dyn. an Off. Publ. Am. Assoc. Anat. 240, 530–565 (2011).

26. Zhou, Q. et al. A multipotent progenitor domain guides pancreatic organogenesis. Dev. Cell 13, 103–114 (2007).

27. Sznurkowska, M. K. et al. Defining Lineage Potential and Fate Behavior of Precursors during Pancreas Development. Dev. Cell 46, 360–375.e5 (2018).

28. Willnow, D. et al. Quantitative lineage analysis identifies a hepato-pancreato-biliary progenitor niche. Nature 597, 87–91 (2021).

29. Escot, S., Willnow, D., Naumann, H., Di Francescantonio, S. & Spagnoli, F. M. Robo signalling controls pancreatic progenitor identity by regulating Tead transcription factors. Nat. Commun. 9, 5082 (2018).

30. Solar, M. et al. Pancreatic exocrine duct cells give rise to insulin-producing beta cells during embryogenesis but not after birth. Dev. Cell 17, 849–860 (2009).

31. Rovira, M. et al. Isolation and characterization of centroacinar/terminal ductal progenitor cells in adult mouse pancreas. Proc. Natl. Acad. Sci. U. S. A. 107, 75– 80 (2010).

32. Single-cell transcriptomics of 20 mouse organs creates a Tabula Muris. Nature 562, 367–372 (2018).

33. Pujal, J. et al. Keratin 7 promoter selectively targets transgene expression to normal and neoplastic pancreatic ductal cells in vitro and in vivo. FASEB J. Off. Publ. Fed. Am. Soc. Exp. Biol. 23, 1366–1375 (2009).

34. Romano, R.-A. et al. ΔNp63 knockout mice reveal its indispensable role as a master regulator of epithelial development and differentiation. Development 139, 772–782 (2012).

35. Backx, E. et al. MECOM permits pancreatic acinar cell dedifferentiation avoiding cell death under stress conditions. Cell Death Differ. (2021) doi:10.1038/s41418-021-00771-6.

36. Xuan, S. et al. Pancreas-specific deletion of mouse Gata4 and Gata6 causes pancreatic agenesis. J. Clin. Invest. 122, 3516–3528 (2012).

37. Szabó, A. & Sahin-Tóth, M. Determinants of chymotrypsin C cleavage specificity in the calcium-binding loop of human cationic trypsinogen. FEBS J. 279, 4283– 4292 (2012).

38. Boros, E. et al. Overlapping Specificity of Duplicated Human Pancreatic Elastase 3 Isoforms and Archetypal Porcine Elastase 1 Provides Clues to Evolution of Digestive Enzymes. J. Biol. Chem. 292, 2690–2702 (2017).

39. Davis, M. A. & Reynolds, A. B. Blocked acinar development, E-cadherin reduction, and intraepithelial neoplasia upon ablation of p120-catenin in the mouse salivary gland. Dev. Cell 10, 21–31 (2006).

40. Hoang, C. Q. et al. Transcriptional Maintenance of Pancreatic Acinar Identity, Differentiation, and Homeostasis by PTF1A. Mol. Cell. Biol. 36, 3033–3047 (2016).

41. Wang, X. et al. Identification of a ΔNp63-Dependent Basal-Like A Subtype-Specific Transcribed Enhancer Program (B-STEP) in Aggressive Pancreatic Ductal Adenocarcinoma. Mol. Cancer Res. 21, 881–891 (2023).

42. Candi, E. et al. Differential roles of p63 isoforms in epidermal development: selective genetic complementation in p63 null mice. Cell Death Differ. 13, 1037– 1047 (2006).

43. Claudinot, S. et al. Tp63-expressing adult epithelial stem cells cross lineages boundaries revealing latent hairy skin competence. Nat. Commun. 11, 5645 (2020).

44. Beer, R. L., Parsons, M. J. & Rovira, M. Centroacinar cells: At the center of pancreas regeneration. Dev. Biol. 413, 8–15 (2016).

45. Seymour, P. A. Sox9: a master regulator of the pancreatic program. Rev. Diabet. Stud. 11, 51–83 (2014).

46. Cleveland, M. H., Sawyer, J. M., Afelik, S., Jensen, J. & Leach, S. D. Exocrine ontogenies: on the development of pancreatic acinar, ductal and centroacinar cells. Semin. Cell Dev. Biol. 23, 711–719 (2012).

47. Mastracci, T. L. & Sussel, L. The endocrine pancreas: insights into development, differentiation, and diabetes. Wiley Interdiscip. Rev. Dev. Biol. 1, 609–628 (2012).

48. Herrera, P. L. et al. Embryogenesis of the murine endocrine pancreas; early expression of pancreatic polypeptide gene. Development 113, 1257–1265 (1991).

49. Kopp, J. L. et al. Sox9+ ductal cells are multipotent progenitors throughout development but do not produce new endocrine cells in the normal or injured adult pancreas. Development 138, 653–665 (2011).

50. Dor, Y., Brown, J., Martinez, O. I. & Melton, D. A. Adult pancreatic beta-cells are formed by self-duplication rather than stem-cell differentiation. Nature 429, 41–46 (2004).

51. Gu, G., Dubauskaite, J. & Melton, D. A. Direct evidence for the pancreatic lineage: NGN3+ cells are islet progenitors and are distinct from duct progenitors. Development 129, 2447–2457 (2002).

52. Backx, E. et al. On the Origin of Pancreatic Cancer: Molecular Tumor Subtypes in Perspective of Exocrine Cell Plasticity. Cell. Mol. Gastroenterol. Hepatol. 13, 1243–1253 (2022).

53. Rooman, I. & Real, F. X. Pancreatic ductal adenocarcinoma and acinar cells: a matter of differentiation and development? Gut 61, 449–458 (2012).

54. Nayak, K. B., Kuila, N., Das Mohapatra, A., Panda, A. K. & Chakraborty, S. EVI1 targets ΔNp63 and upregulates the cyclin dependent kinase inhibitor p21 independent of p53 to delay cell cycle progression and cell proliferation in colon cancer cells. Int. J. Biochem. Cell Biol. 45, 1568–1576 (2013).

55. Nguyen, B.-C. et al. Cross-regulation between Notch and p63 in keratinocyte commitment to differentiation. Genes Dev. 20, 1028–1042 (2006).

56. Patturajan, M. et al. DeltaNp63 induces beta-catenin nuclear accumulation and signaling. Cancer Cell 1, 369–379 (2002).

57. Chari, N. S. et al. Interaction between the TP63 and SHH pathways is an important determinant of epidermal homeostasis. Cell Death Differ. 20, 1080– 1088 (2013).

58. Ripamonti, F. et al. EGFR through STAT3 modulates ΔN63α expression to sustain tumor-initiating cell proliferation in squamous cell carcinomas. J. Cell. Physiol. 228, 871–878 (2013).

59. Maia-Silva, D. et al. Interaction between MED12 and ΔNp63 activates basal identity in pancreatic ductal adenocarcinoma. Nat. Genet. (2024) doi:10.1038/s41588-024-01790-y.

60. Maia-Silva, D. et al. Marker-based CRISPR screening reveals a MED12-p63 interaction that activates basal identity in pancreatic ductal adenocarcinoma. bioRxiv : the preprint server for biology (2023) doi:10.1101/2023.10.24.563848.

61. Fisher, M. L., Balinth, S. & Mills, A. A. ΔNp63α in cancer: importance and therapeutic opportunities. Trends Cell Biol. 33, 280–292 (2023).

62. Ramsey, M. R., He, L., Forster, N., Ory, B. & Ellisen, L. W. Physical association of HDAC1 and HDAC2 with p63 mediates transcriptional repression and tumor maintenance in squamous cell carcinoma. Cancer Res. 71, 4373–4379 (2011).

63. Byrnes, L. E. et al. Lineage dynamics of murine pancreatic development at single-cell resolution. Nat. Commun. 9, 1–17 (2018).

64. Satija, R., Farrell, J. A., Gennert, D., Schier, A. F. & Regev, A. Spatial reconstruction of single-cell gene expression data. Nat. Biotechnol. 33, 495–502 (2015).

65. Dobin, A. et al. STAR: ultrafast universal RNA-seq aligner. Bioinformatics 29, 15– 21 (2013).

66. Anders, S., Pyl, P. T. & Huber, W. HTSeq--a Python framework to work with high-throughput sequencing data. Bioinformatics 31, 166–169 (2015).

67. Love, M. I., Huber, W. & Anders, S. Moderated estimation of fold change and dispersion for RNA-seq data with DESeq2. Genome Biol. 15, 550 (2014).

